# Social preference in rats not impacted by posterior parietal activity despite overall changes in familiarity-based social behavior

**DOI:** 10.1101/2021.11.24.469924

**Authors:** Taylor B. Wise, Rebecca D. Burwell, Victoria L. Templer

## Abstract

Recent literature points to a potential link between the evolution of complex social behavior and the posterior parietal cortex (PPC) in primates including humans (Parkinson & Wheatley, 2013). Thus far, this theory has been overlooked in other highly social animals that may have also evolved due to social selective pressures. In rodents, there is limited knowledge on the involvement of the PPC on sociality, and most studies of such behavior are limited to understanding social preference. We investigated the role of the PPC through two experiments using the 3-Chamber Sociability and Social Novelty test in rats (Crawley, 2004). In Experiment 1, we used a standard 3-Chamber paradigm, which included two novel demonstrators. In Experiment 2, this paradigm was altered to increase the difference in familiarity between demonstrators such that one demonstrator was highly familiar to the subject and the other was entirely novel. Rats with pre-testing permanent neurotoxic lesions were compared to sham surgery control rats, and the same rats were used for both experiments. Experiments 1 and 2 showed that both groups of rats preferred general social interaction, suggesting no deficit in sociability following PPC damage, regardless of demonstrator identity. Further, experimental and control rats showed similar levels of novelty preference following PPC damage, with novelty preferences increasing in Experiment 2. We argue that heightened novelty preference in Experiment 2 may reflect the increased difference in familiarity between demonstrators. Within the confines of the 3-Chamber task, our results suggest that PPC function was not required for general sociability or social novelty recognition. Because the PPC is implicated in abstract cognition, we argue that existing social tests in rodents may not adequately measure the complex cognitive capacities thought to be supported by the PPC. Future studies should investigate the role of the PPC in social cognition by employing behavioral tasks that require higher cognitive demand rather than testing inherent preference for social partners. Outside of our investigation of the PPC, these results show that social novelty preference can be manipulated through changes in familiarity of demonstrators, and that rats can discriminate others’ social identities.

## Introduction

Throughout our evolution selective pressures have molded the brain to manage the intricacies of sociality through complex cognitive systems. Changes in social cognitive function have occurred alongside supporting neural structure and mechanisms, though how these alterations occurred has yet to be fully explained. In particular, many questions on the origins of social cognition are unanswered despite its considerable relevance in our everyday lives. Popular theories such as the social intelligence (Byrne & Bates, 2010), cultural intelligence (Herrmann, Call, Hernández-Lloreda, Hare, & Tomasello, 2007), and the ecological intelligence hypotheses (Rosati, 2017) seek to explain the interaction between social behavior and cognitive evolution from a comparative standpoint. From a neuroscience perspective, others have focused on the evolution of particular neural structures that may reflect social function. One theory posits that the primate brain has been equipped to understand social relationships through a process known as *exaptation*. Through exaptation, a specific region of the brain originally used to manage one aspect of cognition undergoes a cortical repurposing in order to meet new cognitive demands (Anderson, 2010; S. J. Gould, 1991). Specific to social cognition, it is predicted that a key region to be cortically repurposed in response to social needs is the posterior parietal cortex (PPC) (Parkinson & Wheatley, 2013; Yamazaki, Hashimoto, & Iriki, 2009).

The PPC has been investigated across multiple species, with an emphasis on its role in spatial cognition (Kesner & Creem-Regehr, 2013; Mohan, de Haan, Mansvelder, & de Kock, 2018; Nitz, 2006). It has been shown that the PPC is responsible for spatial processing including knowledge on spatial position (Snyder, Batista, & Andersen, 2000), direction of movement and locomotor behavior (Chen, Lin, Green, Barnes, & McNaughton, 1994; Nitz, 2012; Steinmetz, Motter, Duffy, & Mountcastle, 1987), as well as required for tracking route progression (Nitz, 2006; Rodriguez, 2010; Wilber, Clark, Forster, Tatsuno, & McNaughton, 2014). Further, neural activity in the rat PPC provides evidence for a role in spatial cognition (Nitz, 2006, 2012).

A secondary emphasis of research on the PPC has been its link to attentional processes (Colby & Goldberg, 1999; Posner, Walker, Friedrich, & Rafal, 1984; R. L. Reep & Corwin, 2009). Examining the effects of PPC damage in human and non-human primates has shown that it is difficult for subjects to disengage attention from one visual stimulus to engage with another (Farah, Wong, Monheit, & Morrow, 1989; Petersen, Robinson, & Currie, 1989; Posner et al., 1984). This impairment in attentional set shifting has also been shown in rodents where lesioned animals had difficulty abandoning their attention from the initial stimulus dimension in a multi-dimensional learning task (Fox, Barense, & Baxter, 2003). Two subregions, the dorsal (dPPC) and caudal PPC (cPPC), show a functional difference in top-down and bottom-up attentional processes, respectively (Bressler, Tang, Sylvester, Shulman, & Corbetta, 2008). In the primate PPC, it has been shown that the dPPC is specifically implicated in top-down attention used to guide goal-orientated behavior, whereas the cPPC is implicated in bottom-up, stimulus driven attention (Shomstein, 2012). This functional differentiation has recently been confirmed in the rat PPC as well (F.-C. Yang, Dokovna, & Burwell, 2021).

With support from these studies it has been argued that a functional homolog of the PPC exists across human and non-human primates (Campbell & Hodos, 1970). In humans, the PPC consists of the superior parietal lobe (SPL; Brodmann Area 5) and the inferior parietal lobe (IPL; Brodmann Area 7), separated by the intraparietal sulcus. A similar anatomy is found in the primate PPC with a defined SPL and IPL (Cavada & Goldman-Rakic, 1989a, 1989b). Research on this potential homolog has been extended to rodents and considerable work has been dedicated to understanding the rat PPC. In both primates and rats the PPC has been anatomically defined (Kolb & Walkey, 1987; Krieg, 1946; R. L. Reep, Corwin, Hashimoto, & Watson, 1984) including thalamic regions that support the PPC for both species (Baleydier & Mauguiere, 1987; Chandler, King, Corwin, & Reep, 1992; Gutierrez, Cola, Seltzer, & Cusick, 2000; R. Reep, Chandler, King, & Corwin, 1994; F. C. Yang, Jacobson, & Burwell, 2017). Further, scientists have begun to define subregions of the rat PPC that may be functionally analogous to its primate counterpart (Palomero-Gallagher & Zilles, 2015; Pérez-Clausell, 1996; Swanson, 2004; F.-C. Yang et al., 2021). With support from electrophysiology and lesion studies it can be argued that the rat is an appropriate translational model for understanding the primate PPC.

While many studies investigating the PPC have focused on its spatial and attentional functions, there is a subset of literature that has explored the region’s ties to abstract cognition. Abstract cognition can be defined by the ability to represent concepts not directly observed in the physical world (Katz, Wright, & Bodily, 2007), including number, time, and social information. Interesting connections to spatial processing have also been made when investigating abstract cognition. For example, it is argued that time (Bonato, Zorzi, & Umiltà, 2012) and number (Hubbard, Piazza, Pinel, & Dehaene, 2005) may be based on cortical circuitry that supports spatial attention and sensorimotor function, like that of the PPC. Given our ancestors’ selective pressures towards group living, Parkinson & Wheatley (2013) argue that the exaptation theory may be important in understanding how the PPC relates to a third component of abstract cognition, social cognition. This idea is supported by research that shows mechanisms devoted to spatial knowledge are co-opted to process abstract cognition including social cognition (Chiao et al., 2009; Yamakawa, Kanai, Matsumura, & Naito, 2009; Yamazaki et al., 2009). In Yamakawa et al. (2009) subjects undergoing an fMRI task showed similar activation of the PPC when making judgements on physical distance judgements and social compatibility, otherwise known as “social distance” judgments. Combined, this suggests that the neural mechanisms supporting our understanding of physical distance are similar to our perception of social distance and that the PPC may be recruited in both cases.

The social cognition literature mentioned has exclusively studied PPC function in human and non-human primates with no recognition yet of rodent models. Given their excellent spatial navigation and ease in attentional tasks, rats have long been used to study cognitive processes that employ the PPC (Bucci, 2009; Nitz, 2006). Though often overlooked, rats are also highly social animals. In the lab, rats have been shown to prefer social over non-social interactions (Douglas, Varlinskaya, & Spear, 2004; Peartree et al., 2012; Templer, Wise, Dayaw, & Dayaw, 2018; Van Loo, Van de Weerd, Van Zutphen, & Baumans, 2004) and are highly motivated by social reward (Kummer et al., 2011; Thiel, Okun, & Neisewander, 2008; Vanderschuren & Trezza, 2013; Varlinskaya, Spear, & Spear, 1999). In the wild, rats live in large social groups highlighting the ecological relevance of social cognition (Barnett, 2017). It is possible that survival in these groups requires knowledge of complex social systems that may implement regions such as the PPC. **Given that rats exhibit keen spatial perception, inherent social behavior, and ease of neural manipulation, we argue they provide an excellent model for testing the role of the PPC in social cognition**. As a first step in examining hypotheses about the role of the PPC in social cognition, we employed standard experimental lesion approaches in combination with an established test of rodent social behavior. The 3-Chamber Sociability and Social Novelty task measures subjects’ preference for social versus non-social experiences as well as social novelty (Crawley, 2004). Otherwise known as Crawley’s Test, this behavioral paradigm is widely accepted in the literature and has been used to research topics ranging from autism to drug use to aging (Bambini-Junior et al., 2014; G. G. Gould et al., 2012; Jaramillo, Liu, Pettersen, Birnbaum, & Powell, 2014; S. Moy et al., 2004; S. S. Moy et al., 2013; Templer et al., 2018)..

In this study, we explore the role of the PPC as it relates to social behavior in the rat model. To test this, we manipulated dPPC function through permanent neurotoxic lesions and tested subjects on the 3-Chamber Sociability and Social Novelty Test in two experiments. In Experiment 1 we conducted the standardized 3-Chamber task typically used to understand rodents’ social preferences. This includes the use of two non-subject, demonstrator rats as stimuli. In Experiment 2 we manipulated the level of subjects’ familiarity between demonstrator rats to magnify potential differences in novelty preference. Results did not reveal lesion effects in either experiment; however increased difference in familiarity in Experiment 2 did alter both sham and lesioned subjects’ social behavior. Overall, we argue that while the 3-Chamber task appropriately examines social preference, it is incapable of decoding the nuanced cognitive complexities attributed to the PPC. A detailed discussion of this, as well as our findings importance within a larger framework of PPC function, social cognition, and exaptation are addressed. To our knowledge, this study is the first to manipulate PPC function in an investigation of rodent social information processing.

## Methods & Materials

### SUBJECTS

Nineteen adult male Long Evans rats were used as subjects for the purposes of this study (Charles River Laboratories). Four additional male Long Evans rats were used as non-subject demonstrators. All rats were pair-housed and maintained on a diet of 85-90% of their free-feeding weight. Water was never restricted. Prior to testing, all rats were well handled by experimenters and habituated to testing environments. Results from these experiments proceed an initial investigation on PPC activity during an attentional task in an operant chamber. All experiments were conducted in accordance with the NIH guidelines for the care and use of rats in research and was approved by the Institutional Animal Care and Use Committee (IACUC) at Brown University.

### NEURAL MANIPULATION

Experimental rats (n=9) received permanent neurotoxic lesions of the dPPC whereas the control rats (n=10) received sham surgeries. All rats were anesthetized with isoflurane and secured in a stereotaxic frame. Once secured, the incisor bar was adjusted such that bregma and lambda were in the same horizontal plane (±0.2 mm) and rats’ skulls were in a flat position. Craniotomies were made with dental drill and dura was removed to allow insertion of a 28-gauge, 1 mL capacity Hamilton syringe (Sigma-Aldrich) into the PPC (AP: -3.6 to -5.64; DV: -.05 to -1.1) (Olsen & Witter, 2016). Neurotoxic lesions were established through 0.09M NMDA (Tocris Bioscience, Minneapolis, MN) in 0.1M phosphate buffer. NMDA was drawn through a glass pipette and then delivered by pressure injection at 0.1 µL/min for one minute at each of the five bilateral sites. Following injections, the pipette was left in place for three minutes then slowly retracted. For sham surgeries, rats received craniotomies, but had no penetration of cortex. For all surgeries, skin was sutured, and rats were able to recover for at least one week prior to handling and behavioral testing.

### BEHAVIOR

#### EXPERIMENT 1 – STANDARD PARADIGM

All test sessions took place inside a three-room, social interaction chamber (30×45in, Noldus Technologies), known as the 3-Chamber Sociability and Social Novelty Test (Crawley, 2004; Kaidanovich-Beilin, Lipina, Vukobradovic, Roder, & Woodgett, 2011). Rooms were separated by two opaque doors that were removed to initiate the beginning of a test phase. The left and right rooms contained holding cages for demonstrator rats (Figure 1). Live tracking via Ethovision was used during all trials where cumulative sniff duration and frequency of visits to each demonstrator rat were measured. To habituate, subjects could freely explore the entire apparatus for five minutes. No demonstrators or holding cages were present during habituation. If subjects did not display anxious behavior, such as increased grooming, freezing, or defecation, they began Phase 1 approximately 24 hours following habituation. If anxiety was observed, habituation was repeated once a day until behavior was extinguished. Demonstrator rats also received 5-minute habituations to their holding cages prior to testing to ensure that they were comfortable with confinement and would not display anxious behavior that may alter any subject’s approach towards the demonstrator.

**Figure 1.**
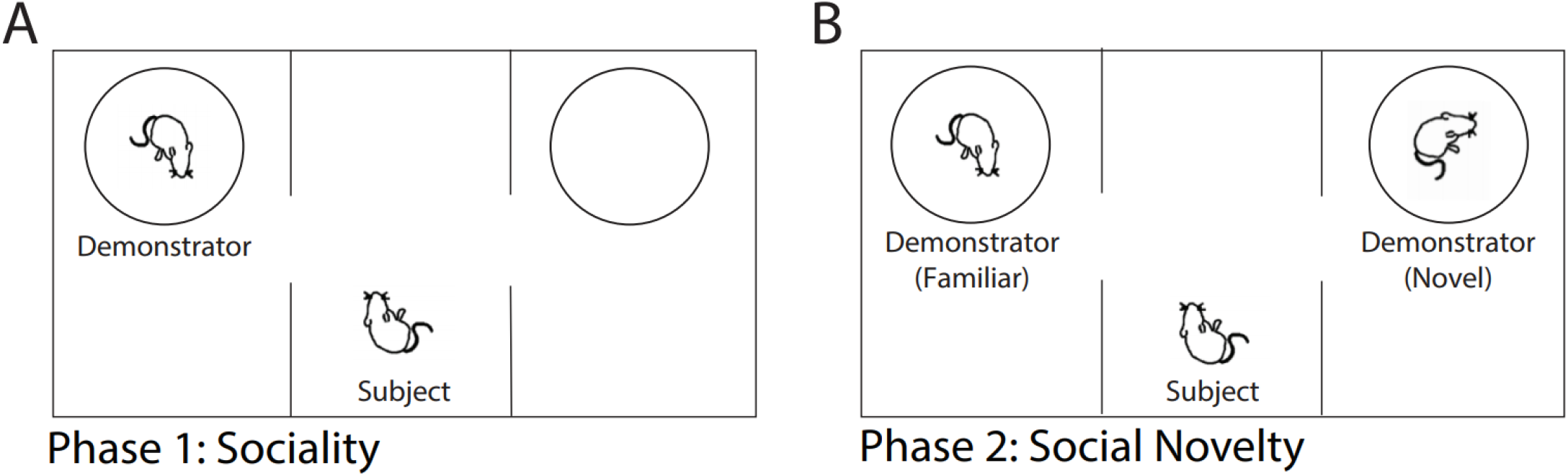
3-Chamber Sociability and Social Novelty task. A) Phase 1-Sociability test in which the subject rat is free to explore a demonstrator rat and an empty cage for 10 minutes. B) Phase 2-Social Novelty test in which a familiar demonstrator rat (continued from Phase 1) is presented along with a novel demonstrator and subject is free to explore for an additional 10 minutes. ITI was approximately 2 minutes between phases.

*Phase 1: Sociability Test*. Round holding cages were placed at one end of the left and right rooms prior to testing (Figure 1A). One cage contained a demonstrator rat unfamiliar to the test subject while the other cage remained empty. The demonstrator/empty cage side was counterbalanced across subjects. To initiate the trial, subjects were placed in the center and doors to the left and right rooms were removed, allowing for exploration of the entire apparatus. The subject could interact with both the novel demonstrator rat and the empty cage as often as desired. Holding cages consisted of evenly spaced bars that allowed demonstrator and subject rats to sniff and see each other while limiting physical contact. Sniff duration to the demonstrator/empty cage was coded by a blind experimenter in real time through Ethovision software. Opaque walls prevented the subject from seeing either cage until entering the respective room. All subjects received a single 10-minute trial and then progressed to Phase 2 of testing.

*Phase 2: Social Novelty Test*. Immediately following Phase 1, subjects were returned to the center room and the doors to either side were closed for an ITI of two minutes. A second unfamiliar rat was placed in the previously empty holding cage while the demonstrator from Phase 1 (now familiar) remained in position (Figure 1B). Designated familiar and novel demonstrators were counterbalanced across subjects. Doors were then removed, and subjects were free to explore for an additional 10 minutes. Identical in form to Phase 1, data was coded by a blind experimenter via Ethovision. Once testing was complete, all rats were returned to their home cages and the apparatus was cleaned to eliminate odor transfer between subjects.

#### EXPERIMENT 2 – INCREASED FAMILIARITY PARADIGM

All methods and analyses were identical to those described in the Experiment 1 apart from introducing subjects to separate demonstrator rats. In the previous experiment, each demonstrator was unfamiliar to the subject prior to testing. In Experiment 2 however, the demonstrator in Phase 1 (thus the familiar demonstrator in Phase 2) was replaced with a cagemate. Subjects were housed with their cagemate for approximately 10 months prior to testing, providing ample opportunity to interact socially. This paradigm was conducted to exaggerate the difference in familiarity between demonstrators in Phase 2, providing an opportunity to examine PPC function in social interaction during varied degrees of presumed familiarity. If the difference in familiarity was exaggerated, we predicted that subjects would show an increased preference towards the novel demonstrator in Phase 2 as compared to their cagemate. Further, if the PPC was required for discriminating highly familiar versus novel social partners, lesions animals would not display this increased novelty preference.

Like in Experiment 1, a habituation process was conducted prior to testing. First, subjects were given a 5-minute habituation in which they could explore the entire apparatus void of any demonstrator rats or holding cages. A second habituation was conducted where subjects could explore their cagemate located inside the holding cage. This served to prevent subjects’ possible anxiety by observing their cagemate in confinement.

##### Phase 1. Sociality Test

Approximately 24 hours after habituation, Phase 1 and Phase 2 were conducted under the same time limits shown in the Standard paradigm. While the prior 5-minute habituation was intended only to diminish potential stress from observing a cagemate in confinement, primary results showed that all subjects preferred their cagemate over the empty cage. These results mirror what would be expected in Phase 1 Sociability Test. In the planned Phase 1, it was shown that social interaction had been nearly extinguished. For this reason, the results we present are from rats’ first experience with their cagemate and will from here forward be referred to as Phase 1, Sociability.

##### Phase 2. Social Novelty Test

All Phase 2 procedures were matched in design to Experiment 1 where subjects were presented with a familiar and novel demonstrator. Critically, in the current experiment, the familiar demonstrator was a *highly* familiar cagemate, while the novel demonstrator remained equally novel as that of Experiment 1. Therefore, subjects’ familiarity to demonstrators across experiments was increased from a 10-minute prior interaction to a 10-month consistent social relationship. Phase 2 consisted of a 10-minute exploration of both demonstrators; however, comparisons across phases were complicated by the shortened 5-minute length of Phase 1 Sociability. To ensure that the 5-minute Phase 1 trial was a true comparison to the 10-minute Phase 2 trial, we analyzed discrimination ratio data from Phase 2 at the 5- and 10-minute mark. In a paired-sample ttest of 5-versus 10-minute data in Phase 2, we found that there was no significant difference in behavior (t=-0.880, p=0.405 for shams; t=-0.431, p=0.678 for lesions). Due to this non-significant difference, we will be comparing the complete time rats spent exploring the apparatus in Phase 1 (5 minutes) to the maximum time of Phase 2 (10 minutes).

#### DATA ANALYSIS

As described above, data was coded blindly and then extracted from Ethovision. Sniff behavior toward the empty cage and all demonstrators was used to calculate a discrimination ratio (DR) where [DR=(Novel-Familiar)/(Novel+Familiar)] (Ennaceur & Delacour, 1988). Discrimination ratios are used to depict complex behavioral data through a single measure, accounting for, in this case, exploration to an empty cage and all demonstrator rats. In our experiments a DR of (+)1 is equal to complete preference for the demonstrator in Phase 1 and for the novel demonstrator in Phase 2, whereas a DR of (-)1 is equal to complete preference for the empty cage in Phase 1 and for the familiar demonstrator in Phase 2. A DR of 0 indicates no preference. Outlier analysis was conducted for all data where outliers were replaced with the mean value of the group and results were analyzed via SPSS.

## Results

### HISTOLOGY

Prior to behavioral testing, subjects received either permanent neurotoxic lesions of the dPPC (n=9) or sham surgeries (n=10). In the experimental group, rats were administered a 0.1 µL NMDA injection at 5 bilateral sites (Olsen & Witter, 2016). A dPPC deficit was achieved with mean percent damage in the left hemisphere at 70% (SD=16.4), with 92% spread (SD=12.2), and a mean percent damage in the right hemisphere at 61% spread (SD=10.1), with 86% spread. Coronal sections illustrating the area affected is shown in Figure 2. Damage outside of the target area was analyzed through percentage of coronal sections displaying bilateral damage to non-PPC cortical regions. Damaged areas included somatosensory regions at 24.1% ±9.6 % and visual cortex at 28.9% ±10.6%. Regardless of outsider cortical damage, no rats showed abnormal behavioral results and thus remained in the study.

**Figure 2.**
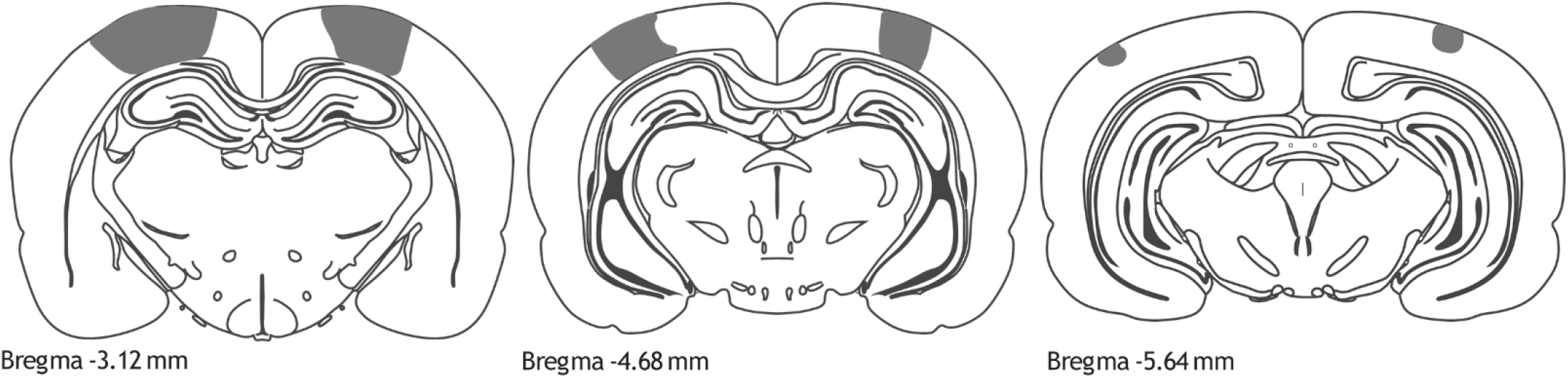
Coronal sections representative areas (shaded in gray) affected from NMDA lesion to dPPC.

### BEHAVIOR

#### EXPERIMENT 1 – STANDARD PARADIGM

In Experiment 1 rats were tested on the standard 3-Chamber Sociability and Social Novelty task. In this paradigm rats were free to explore a novel demonstrator and an empty cage in Phase 1, and a familiar and novel demonstrator in Phase 2. Subjects’ sniff behavior to social stimuli was the primary measure of exploration and converted into discrimination ratios.

For Phase 1, both sham and lesioned rats preferred to spend more time with the demonstrator rat as compared to the empty cage (Figure 3). Discrimination ratios for both groups were significantly different from zero (One-sample ttest: t=4.374, p=0.002 for shams; t=7.654, p=0.000 for lesions) and no group difference in DR scores were shown (Paired-samples ttest: t=-0.545, p=0.600). Together this shows that PPC lesions did not affect rats’ general sociability preferences.

**Figure 3.**
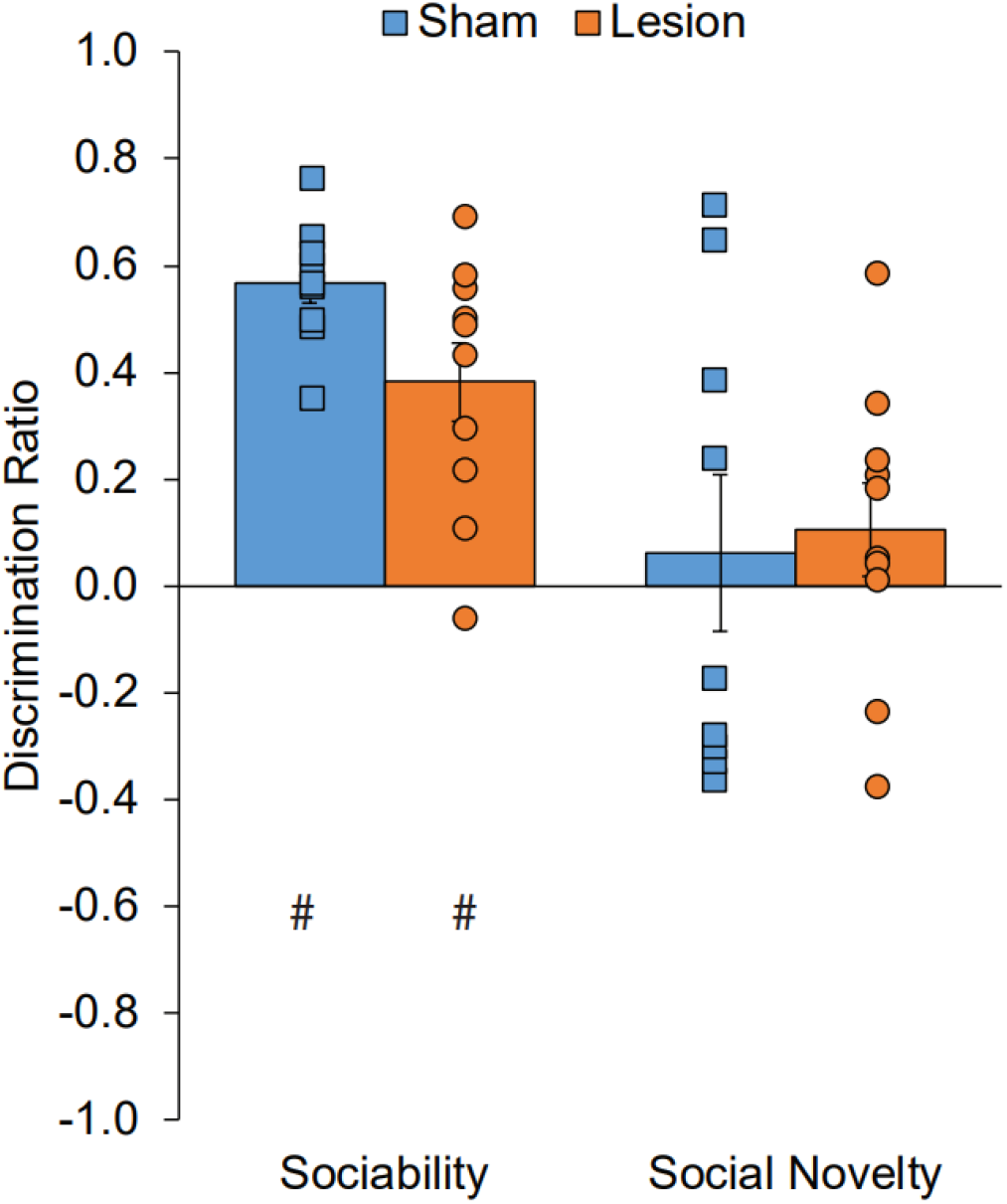
Standard paradigm. Novelty was assessed by a discrimination ratio (DR) indicating the difference in exploration between holding cages as a proportion of the total exploration time. Positive DR indicates preference for social interaction in Phase 1 and social novelty in Phase 2, while a DR of zero indicates no preference. # indicates a significant one-sample t-test (p < 0.05).

In Phase 2, neither group showed a significant preference for either the novel or familiar demonstrator rat, as shown in DR results (One-sample ttest: t=0.891, p=0.443 for shams; t=0.516, p=0.265 for lesions; Figure 3). Again, no group difference in DR scores was shown (Paired-samples ttest: t=0.562, p=0.589). It would be expected that sham animals in this experiment should replicate behavior shown in previous literature, where normal functioning rats prefer social novelty; however, this was not the case in Experiment 1. Possible reasons as to why we did not observe social novelty preference in Experiment 1 are discussed in more depth in our Discussion below.

Our results show that in Phase 1, where rats were free to explore a novel demonstrator and an empty cage, all subjects preferred the social experience. This was true regardless of damage to the PPC, and replicates prior literature on rodent sociability. In Phase 2, when allowed to interact with a novel and familiar demonstrator, no group showed significant novelty preference. Due to sham animals’ nonsignificant DR score we cannot sufficiently claim that lesioned rats’ social novelty preference was affected by the PPC. Importantly, no difference in total exploration was found in either phase when comparing sham and lesioned animals and is thus not thought to be responsible for shams’ decreased social novelty behavior.

#### EXPERIMENT 2 – INCREASED FAMILIARITY PARADIGM

In Experiment 2, rats were again tested on the 3-Chamber task, yet now with a manipulation to increase the difference in familiarity between demonstrator rats. Experimental design was replicated from Experiment 1 with the exception that the demonstrator in Phase 1 (and thus the familiar demonstrator in Phase 2) was replaced with the subject’s cagemate. Because lesioned rats did not prefer social novelty in Experiment 1, substituting the demonstrator with a cagemate may better examine the PPC’s role in social novelty preference. It was hypothesized that by increasing the difference in familiarity between demonstrators in Phase 2, novelty preference would increase, and results may shed light on the level of influence the PPC has over perceived social novelty.

For Phase 1, rats explored their cagemate, still confined in a holding chamber, and an empty cage. Both sham and lesioned rats preferred their cagemate with DR scores significantly different from zero (One-sample ttest: t=2.282, p=0.052 for shams; t=4.758, p=0.001 for lesions) and no group difference (Paired samples ttest: t=-0.778, p=0.459; Figure 4). Replicating results from in Experiment 1, all rats preferred social interaction over an empty cage, even if highly familiar with the demonstrator.

**Figure 4.**
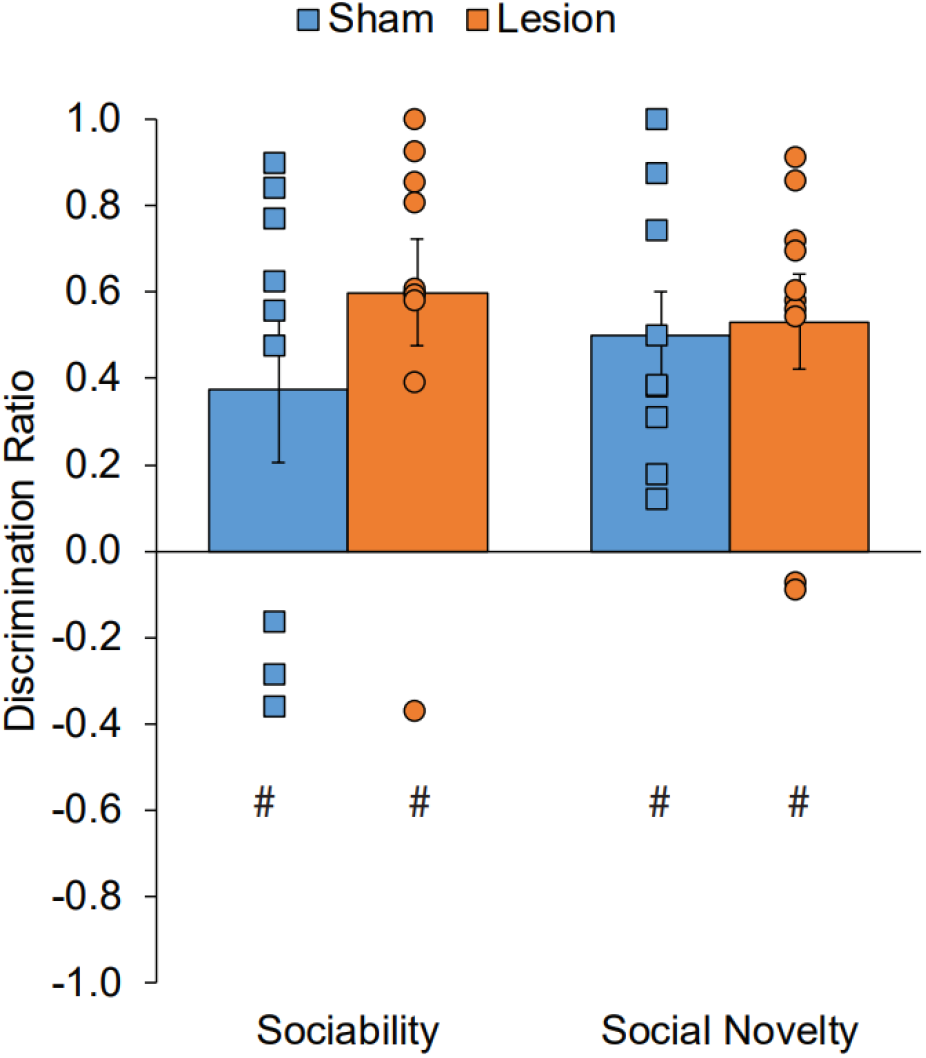
Increased Familiarity paradigm. Novelty was assessed through a DR as described in Experiment 1. # indicates a significant one-sample t-test (p < 0.05). In Phase 1, both groups preferred their cagemate as compared to an empty cage. In Phase 2, both groups preferred social novelty over their familiar cagemate.

In Phase 2, both sham and lesioned subjects preferred to socialize with the novel demonstrator over their cagemate, as shown through DR scores significantly different from zero (One-sample ttest: t=4.205, p=0.003 for shams; t=5.342, p=0.001 for lesions; Figure 4). This suggests that an increased difference in familiarity promoted a heighten novelty preference from Experiment 1. No group difference was found between DR scores (Paired-samples ttest: t=-1.626, p=0.143), suggesting that the PPC did not play a role in social novelty preference. It should also be noted that no group difference in total exploration was found ensuring that results have not been skewed by disparities in overall subject participation.

Overall, Experiment 2 investigated the potential difference in sociability and social novelty preference when a demonstrator rat was replaced with an extremely familiar cagemate. In Phase 1, all rats still preferred social interaction over an empty cage, despite the fact that they had ample opportunity to interact with their cagemate outside of the task. When increasing the difference in familiarity between demonstrator rats in Phase 2, it was found that both sham and lesioned subjects preferred the novel demonstrator over their cagemate. Compared to Experiment 1, lesioned rats now showed a significant novelty preference when a large difference in familiarity was present. Differences in sociability and social novelty between sham and lesioned animals were not observed. Together, these results suggest that regardless of PPC damage, all rats prefer social interactions and prefer demonstrators that are comparably novel.

## Discussion

In this study we examined the hypothesis that the PPC supports social behavior in rats. We tested this hypothesis in two experiments in which we compared social behavior of rats with and without PPC damage, first in a paradigm standard to the literature and second in a paradigm that manipulated the difference in familiarity between non-subject demonstrator rats. Using sniff behavior as a measure of exploration, discrimination ratios were calculated for each experiment to illustrate subjects’ social preferences. By comparing social behavior of rats with and without PPC damage we predicted that, if the PPC plays a role in social preference, lesioned animals would show decreased sociability and decreased social novelty preference in the 3-Chamber task.

In Experiment 1 we conducted the 3-Chamber task with one unfamiliar demonstrator in Phase 1 and a second unfamiliar demonstrator in Phase 2 (Figure 1). In Phase 1, where subjects could explore an unfamiliar demonstrator and an empty cage, both sham and lesioned animals preferred the demonstrator, suggesting that PPC deficits did not affect rats’ affinity for social interaction. This result is fairly unsurprising given that no literature has yet shown that the PPC is required for general sociability, and instead that it is active when making judgements between potential social partners (Chiao et al., 2009; Yamakawa et al., 2009). In Phase 2, where subjects could explore a familiar demonstrator (known from Phase 1) and an unfamiliar demonstrator, both groups showed no preference for either rat. In prior literature it has been shown that normal-functioning rodents exhibit a high social novelty preference (Crawley, 2004; Templer et al., 2018); however, this result was not replicated in our sham animals. There are a few possibilities for these deviated results including an adverse response to the novel demonstrator or heightened interest in the familiar demonstrator. While subjects could never make full-body contact with demonstrators, the holding chamber bars were far enough apart to allow for nose-to-body contact. It has been shown that even when body contact is blocked by barred holding chambers or a mesh wall, rats will respond to the presence of a conspecific (Peartree et al., 2012). Therefore, if specific demonstrators displayed either positive or negative social signals it could elicit a behavioral response from the subject. This may have been the case in Experiment 1, Phase 2. To ensure that these possible confounds did not affect future experiments, new demonstrators were used in Experiment 2. Given this discrepancy with our sham animals we cannot claim that lack of social novelty preference in lesioned rats was due to a PPC deficit.

The 3-Chamber task was again conducted in Experiment 2 with a critical manipulation to the level of familiarity between demonstrator rats. In the Increased Familiarity paradigm, subjects were exposed to a cagemate in Phase 1, and then again in Phase 2 alongside a novel conspecific. Where in Experiment 1 demonstrators only differed in familiarity by the 10 minutes allotted from Phase 1, this experiment examined social novelty preference when a stark difference between the demonstrators’ relationship to the subject existed. To our knowledge this paradigm has yet to be studied and thus provides new insight on how rats respond to highly familiar versus novel social conditions. We hypothesized that when manipulating the difference in familiarity between demonstrators, subjects would alter their social behavior observed from Experiment 1.

In Experiment 2, Phase 1, subjects were first presented with their cagemate and an empty cage. Both sham and lesioned animals preferred their cagemate. Like Experiment 1, this suggests that irrespective of PPC function, rats preferred social over nonsocial experiences. While we did not see PPC effects in Phase 1 for either experiment, these results help to establish what aspects of social behavior are and are not implicated by the PPC. Importantly, all animals in Experiment 2 preferred social over non-social interactions even when their demonstrator was a highly familiar cagemate that they had full-time access to outside of the task. This highlights rats’ strong desire for social stimulation and provides support for the argument that rodent social behavior is not only ecologically relevant, but also worth investigating. In Phase 2, where subjects were presented with their cagemate and a novel demonstrator, both sham and lesioned rats preferred to spend more time in the novel context. Theoretically, this increased difference in demonstrator familiarity should make for a heightened social novelty preference in normal-functioning rats, which we observed. Interestingly we observed this novelty preference in lesioned animals as well. This suggests that the PPC does not play a role in social novelty preference when demonstrator rats exhibit a large difference in familiarity. Alternatively, it is possible that despite PPC dysfunction, the level of novelty was high enough to compensate for any neural deficits. It should also be noted that shams’ social novelty preference in Experiment 2 replicates prior literature, thus supporting our claim that subjects’ unexpectedly low social novelty preference in Experiment 1 may have been due to positive or adverse responses to specific demonstrators rather than inherent social deficits in PPC-lesioned rats.

Overall, both experiments show no group differences between sham and lesioned subjects’ behavior. This is true for both general sociability and social novelty preference. Experiment 2 provides particularly interesting results given its familiarity manipulation. Even with a high familiarity to their cagemate in Phase 1, subjects still chose a social experience rather than a nonsocial one, pointing to rats’ general inclination towards sociality. Further, when manipulating the difference in familiarity, social novelty preference in Phase 2 was altered in both groups. This suggests that PPC damage did not impact novelty preference or recognition. It is possible that this increased difference in familiarity compensated for any PPC damage attributable to low levels of novelty preference; however, it is more likely that the PPC is not implicated in these general social preferences. Regardless of PPC activity, it is possible that if the level of familiarity was to be titrated even further, we would observe a gradient of social novelty preference. A focus on demonstrators’ familiarity to the subject should be included in future studies to better understand the intricacies of rats’ social preferences.

From this study we did not observe any behavioral differences between sham and lesioned animals in terms of sociability or social novelty preference. On the surface this may appear to suggest that the rodent PPC is not implicated in social cognition. However, given that the demands of social cognition are complex and not well understood it can be argued that the 3-Chamber task is simply not equipped to measure these processes. The PPC is considered a multisensory hub where separate streams of information are integrated. Information integration may explain PPC actitivy attributed to abstract cognition including social cognition as well as the evolutionary ties between spatial and abstract domains. The potential role for the PPC in rodent social cognition has yet to be explored in any study outside of the one described here. For this reason, we felt that the investigation must begin with a task widely used and trusted in the field. The benefits of conducting the 3-Chamber task include ease of comparability between our lesioned animals and prior research; however this paradigm does not include many features typically attributed to cognitive tasks such as training, reward, and choice. It also does not require rats to maintain larger connections between multiple streams of information. For these reasons, we argue that the 3-Chamber task does not accurately measure rodent social cognition in relation to the PPC. While we cannot conclude that the PPC is required for rats’ social cognition from our experiments, we can state that this brain region is not implemented in social *preferences*. At the core of the 3-Chamber task, it is a passive test of behavior that never requires rats to learn or maintain extended knowledge of the quality of their social relationships. Of the research that has examined social functions of the PPC thus far, appropriate species specific tasks for measuring PPC activity have involved higher order understandings of social networks and ego-centric bonds. Thus, to better understand possible PPC function in rats it may be necessary to employ behavioral tasks that require high-level cognitive manipulation of social knowledge.

In this study, we took the initial steps required to examine the role of the rodent PPC in social cognition using tasks that are standard in the field. These results are the first report of 1) PPC function in the 3-Chamber test, 2) the PPC-social cognition link in rodents, and 3) social novelty manipulation, thus adding value to our limited knowledge on rodent social cognition and PPC activity. Although we did not observed effects of PPC damage on the 3-Chamber Sociability and Social Novelty task; we did find that social novelty preference was increased following a manipulation to the level of familiarity between demonstrators. Our findings regarding the role of the PPC in social cognition may be due to the simplicity of the task and the reliance on innate preference behaviors. In future experiments we plan to examine the potential mechanistic overlap of spatial and social cognition using novel tasks that require more complex manipulation of social information.

## Notes

### Competing Interest Statement

The authors have declared no competing interest.

